# Afferents to Action: Cortical proprioceptive processing assessed with corticokinematic coherence specifically predicts gross motor skills

**DOI:** 10.1101/2023.09.26.559273

**Authors:** Mongold Scott, Georgiev Christian, Legrand Thomas, Bourguignon Mathieu

## Abstract

Voluntary motor control is thought to be predicated on the ability to efficiently integrate and process somatosensory afferent information. However, current approaches in the field of motor control have not factored in objective markers of how the brain actually tracks incoming somatosensory information. Here, we asked whether motor performance relates with such markers obtained with an analysis of the coupling between peripheral kinematics and cortical oscillations during continuous movements, best known as corticokinematic coherence (CKC). Motor performance was evaluated by measuring both gross and fine motor skills using the Box and Blocks Test (BBT) and the Purdue Pegboard Test (PPT), respectively, and with a biomechanics measure of coordination. Sixty-one participants completed the BBT, while equipped with electroencephalography and electromyography, and the PPT. We evaluated CKC, from the signals collected during the BBT, as the coherence between movement rhythmicity and brain activity, and coordination as the cross-correlation between muscle activity. CKC at movements’ first harmonic was positively associated with BBT scores, and showed a relationship with PPT scores, but only in synergy with BBT scores, where participants with lower PPT score had higher CKC than expected based on their BBT score. Coordination was not associated with motor performance and at most, weakly related to CKC. These findings demonstrate that cortical somatosensory processing in the form of strengthened brain-peripheral coupling is specifically associated with better gross motor skills. CKC might be considered as a valuable addition to classical tests of proprioceptive acuity, with important perspectives for future clinical studies and practice.

**Significance Statement:** Whether standing upright, jogging, or in Olympic competition, our nervous system not only sends out motor commands prompting muscles to contract, but also receives incoming information to fine-tune motor actions. Though the machinery involved in sensing mechanical changes is well-described, the neural processing of this information is not, making its relevance to motor function unresolved. We found that the coupling strength between peripheral kinematics and cortical activity was related to motor function and at most, only weakly related to conventional muscle-only assessments. We present novel behavioral relevance of this coupling and its specific relationship to gross motor skill. Our study paves the way for including novel brain-centered approaches to complement classical assessment sensorimotor functions in health and disease.

## Introduction

Proprioception is the sense of position and movement of our own body. While the ‘hardware’ of proprioception, consisting of a collection of sensory mechanoreceptors located in muscles, tendons, joints, and the skin, provides kinematic data for the brain, central processing, akin to ‘software,’ must integrate and use this information to update motor programs (Han et al., 2016; Imai and Yoshida, 2018; Palva and Palva, 2018; Tuthill and Azim, 2018). Currently, much of the literature describing the proprioceptive sense relies on peripheral proprioceptive acuity testing, with common tasks including threshold to detection of passive motion, joint position reproduction, or active movement extent discrimination. While these tasks have been useful in risk identification or in a variety of clinical and sport-specific settings (Noronha et al., 2006; Hobbs et al., 2010; Mir et al., 2014), they do not describe the ‘software’ of proprioception. And although the cortical and subcortical areas active during proprioceptive stimulation are well known (Goble et al., 2011), the neurophysiological mechanisms therein are still poorly described. The characterization of these mechanisms appears crucial considering that changes in movement sense are implicated across countless pathologies (Dietz, 2002), during aging (Ferlinc et al., 2019), and in competitive sport (Han et al., 2016).

It is clear that the field of motor control would benefit by an approach that examines how the brain tracks incoming proprioceptive information. Correspondingly, corticokinematic coherence (CKC) is a neurophysiological method that is thought to specifically assess the cortical processing of somatosensory proprioceptive afferences (Piitulainen et al., 2013a, 2018a; Bourguignon et al., 2015). CKC analyses during repetitive movement tasks have revealed a significant coupling between oscillatory cortical activity and limb kinematics at movement frequencies (Bourguignon et al., 2011, 2012; Piitulainen et al., 2013a). They have already revealed differences in proprioception in the case of children with cerebral palsy and in elderly adults (Piitulainen et al., 2018b; Démas et al., 2022). Though CKC appears to be a viable tool to probe the neural tracking of proprioceptive information, its relationship to motor performance is yet unexplored. Motor performance is a highly multidimensional concept. A common subdivision distinguishes between gross and fine motor skills. In this regard, the Box and Blocks Test (BBT) and the Purdue Pegboard Test (PPT) are regularly used in the rehabilitative medicine field to characterize these two aspects of motor performance. While both tasks require use of the upper extremity, the BBT relies more on gross motor function, whereas the PPT necessitates more fine motor control (Gardner and Broman, 1979; Mathiowetz et al., 1985). Importantly, the highly repetitive nature of the BBT makes it an especially suitable task for CKC analysis (Marty et al., 2015a).

Yet another determinant contributor to motor performance is coordination (Bernshteĭn, 1967). In biomechanics, coordination is defined as the relation between muscle activations acting toward the achievement of a motor task (Kimura et al., 2021). The neuronal underpinning of coordination is not fully understood, but an increasingly acknowledged mechanism is the simultaneous control of multiple muscles by specialized spinal circuits, commonly known as muscle synergies (Saltiel et al., 2001; d’Avella et al., 2003; Kuppuswamy and Harris, 2014). Motor coordination can be assessed through the electromyographic (EMG) assessment of neuromuscular activations between muscles.

In this study, we estimated CKC with the aim of assessing its relationship to two types of motor skills, as measured by the BBT (gross) and PPT (fine). Additionally, coordination during the BBT was assessed to test its relationship with motor performance and with CKC. Correspondingly, we hypothesized that *(i)* CKC strength would be positively correlated with BBT and PPT scores, and that *(ii)* enhanced muscular coordination would coincide with better motor scores. Finally, since CKC is thought to represent the processing of incoming sensory afferents, and coordination to rely on spinal mechanisms for motor control, we hypothesized that *(iii)* these two measures would be at most, weakly related.

## Methods

### Participants

Sixty one healthy adults (32 females; mean ± SD age, 23.9 ± 3.5 years, range 18–34 years) participated in the study. All participants were right-handed according to the Edinburgh Handedness Inventory (mean ± SD score, 72.6 ± 9.9; range, 57 – 90; Oldfield, 1971) and reported no history of neurological or motor impairments. Participants had no prior experience with the BBT and PPT. The protocol was approved by the ethics committee of the Université Libre de Bruxelles. All subjects gave written informed consent in accordance with the Declaration of Helsinki.

### Experimental Design

Participants were fitted with an EEG cap with 64 electrodes (Waveguard original, ANT Neuro). Additionally, participants were fitted with electrodes on three muscles of the right upper extremity: first dorsal interosseous (FDI), biceps brachii, and anterior deltoid. Participants completed three trials of the BBT with the right hand. The BBT requires a participant to transfer as many small wooden blocks as possible across a 15.2-cm tall barrier with the designated testing hand, during one minute of testing (Mathiowetz et al., 1985). One hundred and fifty 2.5-cm^3^ wooden blocks were placed in random orientations in the partition on the side of the testing hand. A participant’s score was equal to the number of blocks moved over the barrier, to an identical, but empty partition, in one minute. The experimenter instructed the participant to move the blocks as quickly as possible and encouraged them throughout each trial.

Participants completed five trials of one subset of the PPT, in which they used only their right hand to transfer metal pegs from their starting position, in the upper right-hand side of the board, to the right column with small holes spaced along the length of the board. Each participant was instructed to transfer as many pegs as possible in 30 seconds.

The experimenter performed one trial of the BBT and PPT to show the participant how to perform it properly.

### Data Acquisition

EEG and EMG signals were recorded during the BBT trials. The EEG signals were continuously recorded with a conventional cap in a layout based on the extended international 10–20 system for electrode placement. The nasion, inion, and the mid-sagittal plane of the head were used as anatomical landmarks to place the cap. The EEG cap was connected to an EEG mobile amplifier (eegoTM, ANT Neuro, Netherlands). Electrode-skin impedance levels were kept below 20 kΩ with electrolyte gel. Brain activity was visualized and sampled at 1000 Hz using the eegoTM software (v1.9.1; ANT Neuro, Netherlands) running on an interactive tablet.

The EMG signals were continuously recorded with bipolar electrodes. EMG electrodes were connected directly to the EEG amplifier and signals were recorded on auxiliary channels of the EEG amplifier. EMG signals were sampled at 1000 Hz. Proper placement of EMG electrodes on respective muscle bellies was verified during isometric contractions.

### Data pre-processing

EEG signals were imported into Matlab (R2019b, MathWorks, Massachusetts, USA) where custom scripts combined with functions from the FieldTrip toolbox (Donders Institute for Brain Cognition and Behavior, Nijmegen, The Netherlands; Oostenveld et al., 2011) were used for data analysis. EEG signals at electrodes affected by excessive noise level (mainly due to bad electrode-skin contact) were reconstructed by interpolation of the signals from the surrounding electrodes as done previously (Perrin et al., 1989). Electrodes were considered noisy when they matched at least one of the three criteria: 1) too high wide-band amplitude, 2) too high ratio between high and low frequency amplitudes, and 3) too low correlation with other channels, as proposed previously (Bigdely-Shamlo et al., 2015). Details of the methods and default values used for these three criteria can be found in Bigdely-Shamlo et al. (2015). EEG signals were then re-referenced to a common average. Thereafter, independent component analysis was applied to the EEG signals filtered through 0.5–45 Hz, with the EEG data being decomposed into 20 independent components using FastICA algorithm (dimension reduction, 25; non-linearity, tanh; Vigario et al., 2000; Hyvärinen et al., 2001). The spatial distribution and time-series of each component were visually analyzed for each trial, with 1–6 components corresponding to eye-blink and heartbeat artifacts, or non-uniform noise selected for removal. The corresponding components were subsequently subtracted from raw EEG signals.

As a preliminary step to correct for slow drift, movement, and power-line artifacts, EMG signals recorded during the BBT trials were filtered between 20 and 295 Hz with notch filters at 50 Hz and harmonics using a zero-lag FFT-based filter and rectified (Fig. 1A). We then applied a processing procedure to correct for changes in EMG amplitude across cycles. To do this, fast and slow envelopes were extracted from each rectified EMG by smoothing it with a gaussian kernel that implements a low-pass filter at 3 Hz and 0.7 Hz, respectively (Fig. 1A). These frequencies were chosen to capture the range of expected movement frequencies (1–1.5 Hz). The ratio of the fast envelope and the slow envelope represents the muscle recruitment trace (Fig. 1B). It emphasizes the periodicity of the individual muscle activity within each movement cycle while removing the influence of non-stationarity across cycles.

**Figure 1.**
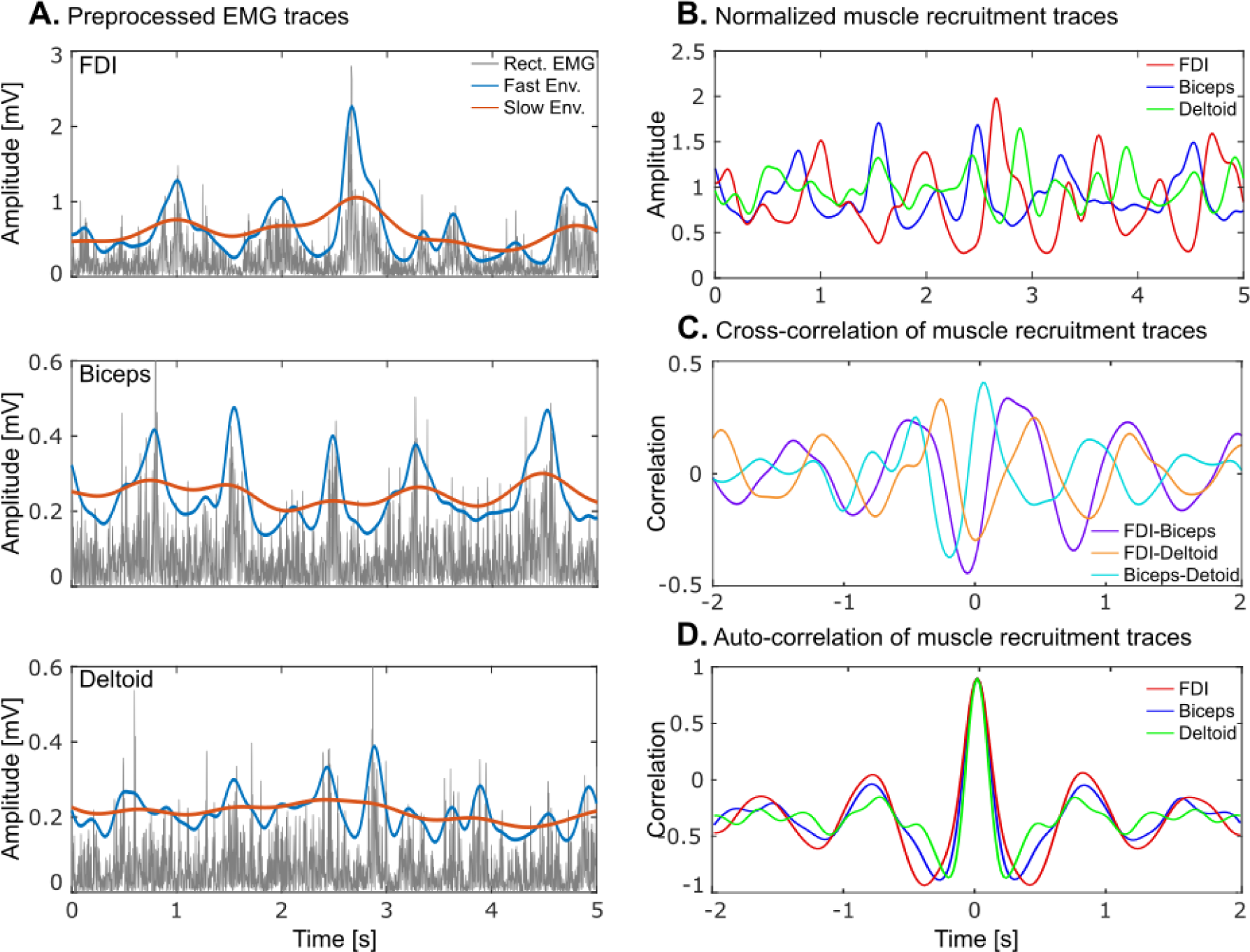
Processing of EMG signals. A — Excerpt of pre-processed EMG signals for the FDI, biceps and deltoid from a representative participant. From each of the rectified EMG signals (dark gray traces) were extracted a fast (blue traces) and a slow (red traces) envelope. B — Muscle recruitment traces for each of the muscles, obtained as the ratio between the fast and slow envelopes. C — Cross-correlations between pairs of muscle recruitment traces. Their maximal amplitude was taken as an estimate of motor coordination. D — Auto correlations for the same muscle recruitment traces. The amplitude of their side peaks was taken as a measure of regularity.

### EMG processing

During the BBT, three parameters of muscle activity were extracted from the upper extremity: coordination, regularity, and EMG modulation depth.

Motor coordination was assessed with two different methods: cross-correlation and mutual information. Both are fit for clinical translation as neither are computationally demanding. Cross-correlation is an established method for comparing EMG signals between different muscles (Hawkes et al., 2012). This approach measures the similarity in shape between two curves, with regard to the temporal characteristics of the signals (Hawkes et al., 2019). Here, the Pearson correlation coefficient was estimated between each pair of the three muscle recruitment traces with delays ranging from −2 to 2 seconds, by steps of 20 ms (Fig. 1C). For each pair, the maximum in absolute value was retained and the average across pairs was taken as the correlation-based coordination measure. Mutual information analysis has been established as another similarity index, with successful use assessing motor coordination (Samani et al., 2017). Importantly, this method detects linear and nonlinear statistical dependencies between time series, whereas cross-correlation is limited to the assessment of linear dependence (Jeong et al., 2001). Upon analysis of the activity of the deltoid muscle across participants, we noticed a biphasic activation pattern along the movement cycle. Therefore, the mutual information approach was adopted to capture potential non-linear relationships between signals that otherwise would not have been identified. Mutual information was estimated between each pair of the three upper extremity muscle recruitment traces. Three types of mutual information were computed and averaged across muscle pairs: (i) mutual information directly applied to the muscle recruitment traces, (ii) mutual information applied to the phase of the muscle recruitment traces, as extracted with the Hilbert transformation, and (iii) permutation mutual information (Cheng et al., 2018). In the two first cases (i and ii), mutual information was applied to the data quantized into 10 equally populated bins. In the last case (iii), mutual information was applied to the permutation order derived from four adjacent time points with a distance of 100 ms. Thus, permutation mutual information determined whether the order obtained by sorting the values over the four time points in one signal tended to predict the presence of one or more other orders in the other signal.

Regularity was assessed because it could bias our estimation of coordination and CKC, the latter two being expected to increase with more regular movements. Regularity can be defined as the consistency of a given muscle activation pattern, where high regularity corresponds to similar activation patterns across cycles of a given task and low regularity results from heterogeneous activation patterns. With this view in mind, regularity can be assessed using autocorrelation, as done more commonly in gait kinematic investigations (Kobayashi et al., 2014) and in studies of sensorimotor coordination (Carroll et al., 2001), where the EMG signal for a designated muscle is compared to itself across successive time lags. The auto-correlation coefficient is 1 at time 0 and decreases with increasing delays imposed on the signal. However, in rhythmic activities, positive side-peaks appear at integer multiples of cycle duration with negative side-peaks in between; the more regular the signal, the greater the magnitude of the side-peaks. Therefore, we estimated the Pearson auto-correlation for three muscle recruitment traces with delays ranging from −2 to 2 seconds, by steps of 20 ms, and quantified regularity with the magnitude in absolute value of the first negative and positive side-peaks (Fig. 1D).

The modulation depth of muscle recruitment traces was assessed to quantify the salience of rhythmic EMG activity. It was quantified with the coefficient of variation (CV), later averaged across the 3 muscles. This parameter is important to control for as it has the potential to bias regularity, coordination, and CKC measures. Indeed, the latter three measures are expected to decrease with decreasing modulation depth, simply because movement cycles become less clearly represented.

### EEG processing

CKC was calculated for the three 1-minute-long trials of the BBT combined. The coherence was computed between rectified FDI muscle activity, which faithfully picked up movement rhythmicity (Piitulainen et al., 2013b; Bourguignon et al., 2019), and each EEG signal. Coherence is an extension of Pearson correlation coefficient to the frequency domain, which quantifies the degree of coupling between two signals, i.e. the CKC strength, by providing a number between 0 (no linear dependency) and 1 (perfect linear dependency) for each frequency (Halliday et al., 1995).

For coherence analyses, the continuous data were split into 5-s epochs with 4-s epoch overlap, leading to a frequency resolution of 0.2 Hz (Bortel and Sovka, 2007). EEG epochs where signal amplitude of at least one electrode exceeded five standard deviations above the mean were excluded to avoid contamination of the data by internal or external noise sources. We then performed coherence analysis (Halliday et al., 1995), yielding cross-, power- and coherence spectra—between EEG and rectified EMG signals. The magnitude squared coherence was chosen as the coupling measure as done in previous CKC studies (Bourguignon et al., 2011, 2012, 2013, 2015; Piitulainen et al., 2013b, 2015; Marty et al., 2015b). Individual power and coherence spectra were used to identify peak CKC, which occurred at the approximate movement frequency (F0) and its first harmonic (F1). Maximum values at F0 and F1 across all electrodes, excluding those located on the edge of the cap, were extracted for each individual. Topographic distributions of CKC, visualized using the FieldTrip toolbox (Halliday et al., 1995; Oostenveld et al., 2011), confirmed that peaks in coherence were identified in the neighborhood of electrodes above the primary sensorimotor cortex (SM1), contralateral to the movement.

The SNR of the EEG signal is a potential confounding factor in coherence analysis, where decreased SNR leads to decreased coherence. To take this factor into account, we assessed it at F0 and F1, for the electrode with maximum CKC. It was estimated as the power at the considered frequency (F0 or F1) divided by the geometric mean of the power ± 2 frequency bins away.

### Statistical analysis

Individual CKC values were compared against a surrogate distribution to determine significance. Surrogate coherence was evaluated between original EEG and Fourier transform surrogate reference signals (1000 repetitions). The Fourier transform surrogate of a signal is obtained by computing its Fourier transform, replacing the phase of the Fourier coefficients by random numbers in the range [-π; π], and then computing the inverse Fourier transform. The threshold for statistical significance was set to *p* < 0.05 and was obtained as the 95^th^ percentile of the distribution of the maximum coherence across 1 – 4 Hz and across all considered electrodes.

Correlation between motor performance scores (BBT and PPT), EMG-based parameters and CKC values were assessed via the Spearman’s rank-order correlation test. It should be noted that a very strong correlation (*r* > 0.8) was identified within the 4 measures of motor coordination and within the 2 measures of regularity. Therefore, related parameters were combined into a unique measure by averaging them after classical standard normalization (mean subtraction and division by the standard deviation).

Potential confounding factors for regularity, coordination and CKC were assessed by correlation and corrected for with linear regression when the correlation was statistically significant. For regularity, the confounding factor was modulation depth as clearer variations in the EMG envelope may correspond to increased regularity. For coordination they were regularity and modulation depth. And for CKC, they were modulation depth, regularity, and EEG SNR.

A partial information decomposition (PID; Ince, 2017; Ince et al., 2017) analysis was used to dissect the information about CKC brought by motor performance. PID is a method to describe the extent by which two explanatory variables bring unique information about a target (information present in one variable but not in the other), redundant information (information common to the two variables), and synergistic information (information emerging from the interaction of the two variables) (Kunert-Graf et al., 2020). In this way, PID was used to estimate the unique, redundant, and synergistic information brought by BBT and PPT performance about CKC. PID analysis was implemented in Matlab, from all three variables with Gaussian copula rank normalization done (Williams and Beer, 2010; Ince et al., 2017; Park et al., 2018). Following, partial information was calculated with a redundancy lattice as done previously (Williams and Beer, 2010; Ince et al., 2017; Park et al., 2018). For statistical assessment and conversion into easily interpretable z-scores, measures of information were compared to the distribution of these measures obtained after permuting motor scores across subjects (1000 permutations).

## Results

### Gross and fine motor skill assessment

Participants’ scores on the BBT, which assesses gross motor skills, ranged from 54 to 98 blocks transported during a single 1-min trial, with an average (± SD) of 76.2 (± 7.6) blocks. These scores are compatible with existing normative data of healthy adults, as reported in females, 78 (± 10.4), and males, 77 (± 11.6) (Mathiowetz et al., 1985).

On the more dextrous task, the PPT, participants’ scores ranged from 11 to 19 inserted pegs, with an average of 15.7 (± 1.7) pegs inserted. These scores are comparable to normative data from young, healthy adults as reported in females, 16.6 (± 2.1), and males, 15.7 (± 1.7) (Yeudall et al., 1986).

Figure 2 presents the relation between BBT and PPT scores. Both were significantly correlated (*r* = 0.39, *p* = 0.002). However, the magnitude of this association was not particularly strong, confirming that BBT and PPT assess partially different aspects of motor performance. Overall, existing variability and decoupling in test scores lends support to the notion that participants possess differing motor control profiles.

**Figure 2.**
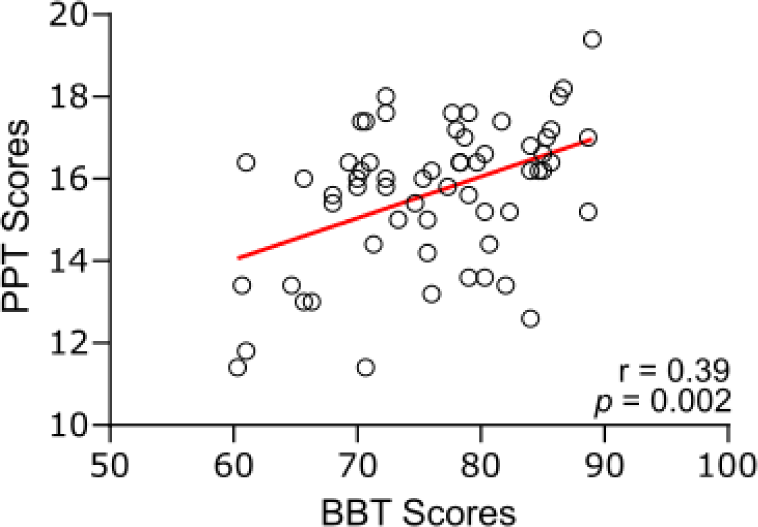
Relationship between PPT scores (pegs inserted in 30 seconds) and BBT scores (blocks transferred in one minute). Circles indicate individual values, and their linear regression line is in red. Correlation value and associated significance level are indicated in the bottom right corner.

### EMG modulation depth, regularity and coordination

The EMG modulation depth across participants possessed a mean (0.23) that was relatively high compared to its SD (0.03), indicating that the modulation of muscle activity was captured similarly across participants.

The two measures of regularity, including the negative and positive auto-correlation side-peaks, were highly related (*r* = 0.81, *p* < 0.0001). Therefore, these two measures were combined into one measure of regularity of muscle activity. This resulting measure of regularity showed a significant positive correlation with EMG modulation depth (*r* = 0.49, *p* < 0.0001). To account for this association, the regularity measure was corrected for EMG modulation depth.

The four coordination measures extracted from the cross correlation and mutual information analyses were highly related (0.80 < *r* < 0.95, *p* < 0.0001). Therefore, these measures were combined into a unique measure of coordination. The resulting coordination measure was significantly correlated with EMG modulation depth (*r* = 0.59, *p* < 0.0001). Consequently, the coordination measure was corrected for EMG modulation depth. In addition, we identified another significant relationship between corrected coordination and corrected regularity measures (*r* = 0.76, *p* < 0.001). Therefore, the coordination measure was corrected for both EMG modulation depth and regularity.

Notably, there was no significant correlation between regularity and BBT (*r* = 0.15, *p* = 0.24) or PPT scores (*r* = 0.12, *p* = 0.37), nor between coordination and BBT (*r* = −0.01, *p* = 0.92) or PPT scores (*r* = 0.07, *p* = 0.62).

### CKC

Figure 3 presents a characteristic coherence spectrum and topography found for the coupling between cortical activity and rectified FDI muscle activity during the BBT. In line with previous studies (Bourguignon et al., 2011, 2012, Piitulainen et al., 2013a, 2018a, 2020; Marty et al., 2015a), such CKC peaked at F0 and F1. Also, the topography at each frequency showed a tendency for CKC to peak approximately around the C3 electrode, compatible with SM1 activation, with sources at times radially or tangentially oriented. Overall, 31 participants showed significant CKC at F0, representing 52% of our sample, while 41 participants had significant CKC at F1, accounting for 67% of our sample. Interestingly, it was not uncommon for CKC to be higher at F1 (0.23 ± 0.10) compared to F0 (0.20 ± 0.11), but this difference was not significant (*p* = 0.08; Wilcoxon test).

**Figure 3.**
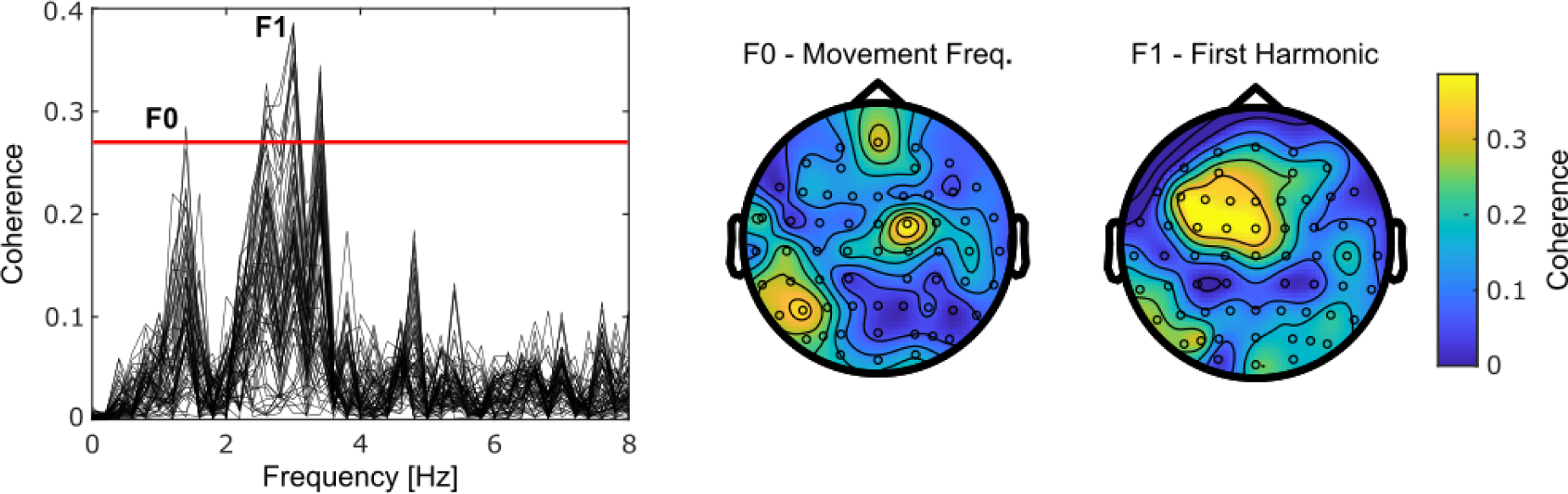
Spectrum and scalp distribution of CKC for a representative individual. The CKC spectrum presents one trace for each EEG electrode, and a horizontal red line indicates the level of statistical significance. Coherence peaked at F0 and F1, corresponding to ∼1.5 and 3 Hz, respectively, for this particular individual. Scalp distributions are mostly compatible with tangential sources in the left SM1 cortex, although other sources might have contributed (Bourguignon et al., 2012).

### Behavioral relevance of CKC

Correlations of CKC at F0 and F1 with the BBT and PPT scores yielded different results. At F0, CKC was not significantly associated with BBT scores (*r* = −0.02, *p* = 0.88) or PPT scores (*r* = −0.11, *p* = 0.40). However, at F1, CKC was significantly associated with BBT scores (*r* = 0.41, *p* = 0.001; Fig. 4A) but not with PPT scores (*r* = 0.07, *p* = 0.60; Fig. 4B).

**Figure 4.**
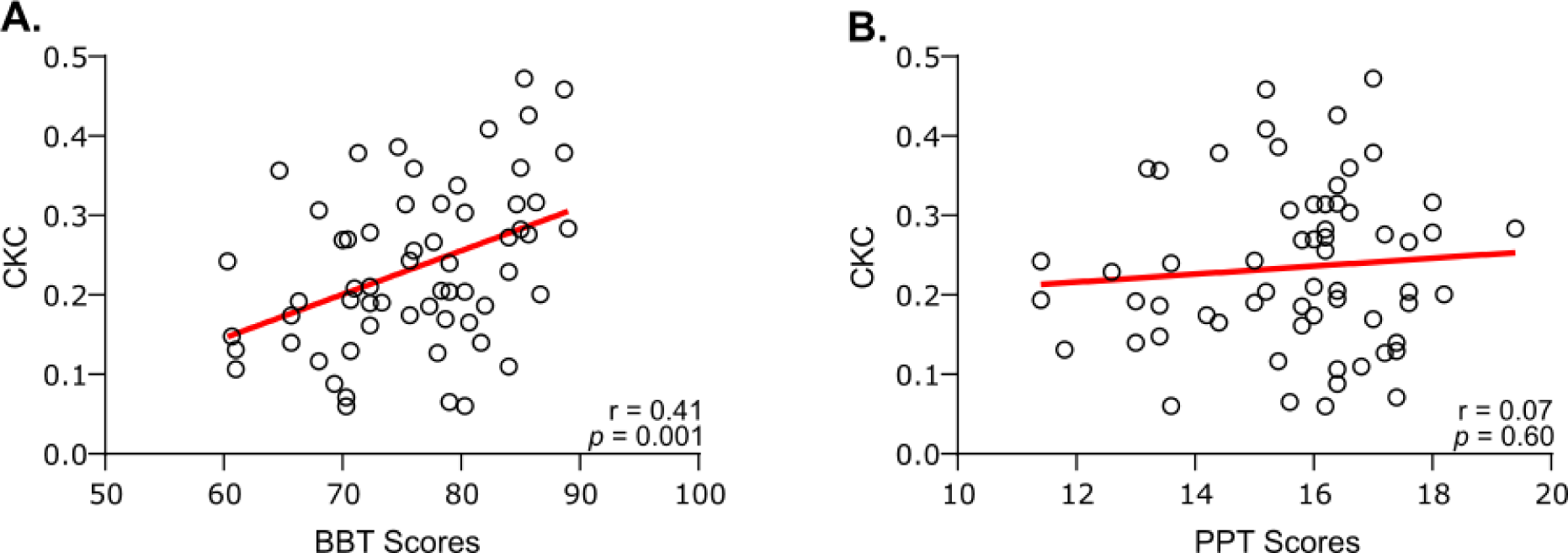
Relation of CKC at F1 with BBT scores (A) and PPT scores (B). Plots are as in Fig. 2.

Additionally, we investigated whether the association between CKC at F1 and performance could be explained by confounding factors. As a result, CKC was not significantly associated with EMG modulation depth (*r* = 0.002, *p* = 0.99), or regularity (*r* = −0.009, *p* = 0.94). These results indicate that the EMG SNR was high enough in all individuals not to noticeably influence CKC values, and that the range of movement regularity was restricted enough not to smear CKC peaks in some individuals more than in others. Our last step to deal with potential confounding factors involved the inclusion of the SNR from EEG activity. CKC at F1 was significantly associated with the SNR (*r* = 0.30, *p* = 0.020), and hence corrected for it. After correction, CKC at F1 was still significantly associated with BBT scores (*r* = 0.29, *p* = 0.021).

Finally, we tested our hypothesis that CKC is at most weakly related to motor coordination. Our data was in line with this hypothesis, revealing non-significant correlation between coordination and CKC at F0 (*r* = 0.24, *p* = 0.07) and F1 (*r* = 0.16, *p* = 0.24). And when CKC data was corrected for EEG SNR, there was a significant correlation at F0 (*r* = 0.29, *p* = 0.02), but not F1 (*r* = 0.20, *p* = 0.12).

### PID

Figure 5A represents the information structure of BBT scores and PPT scores as related to the prediction of CKC. BBT scores brought significant unique information about CKC (*z* = 9.12; *p* = 0.002), whereas PPT scores did not (*z* = −0.98; *p* = 0.97). Together, the two scores did not bring significant redundant information (*z* = 0.38; *p* = 0.20), but their simultaneous observation did bring synergistic information about CKC (*z* = 2.58; *p* = 0.03).

**Figure 5.**
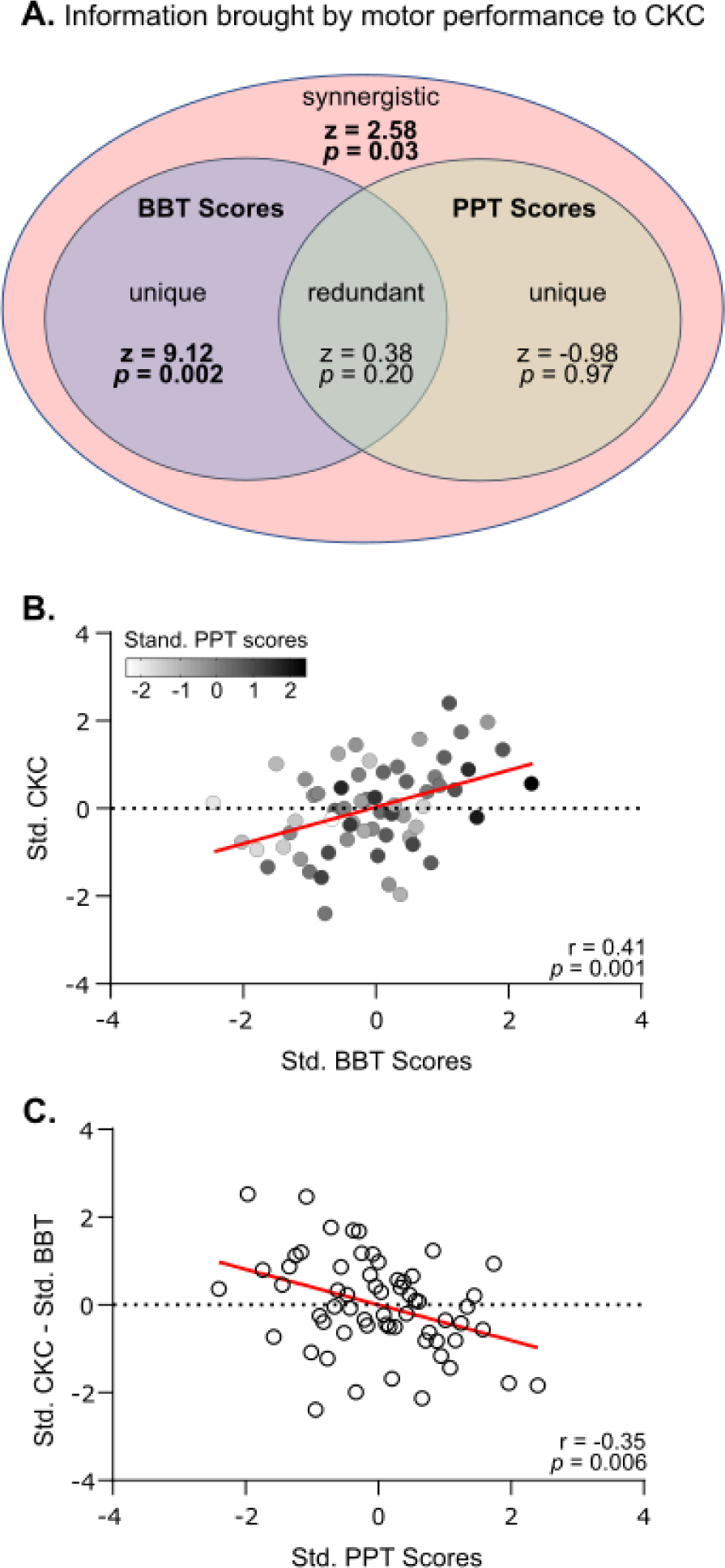
Fine-grained relationship of CKC with motor skills. A — Information structure of BBT and PPT scores predicting CKC. Ellipses outline the decomposition of mutual information into unique information brought by BBT scores (purple), unique information brought by PPT scores (beige), redundant information present in both scores, and synergistic information not present in either scores but emerging from them together. Their significance level is indicated in each partition. B — Relationship between standardized CKC and standardized BBT scores, where individual values are color-coded for their standardized PPT scores. C — Relationship between standardized BBT scores subtracted from standardized CKC as a function of PPT scores. Plots are as in Fig. 2.

Figure 5B characterizes how CKC varies according to BBT and PPT scores. Clearly, from the correlation analysis, CKC is best predicted by BBT scores compared to PPT scores, and this is supported by the difference in unique information that each brings to CKC. However, when plotting the variables together (Fig. 5B), it becomes apparent that the knowledge of the PPT scores could help refine CKC prediction beyond its regression line on BBT scores. Participants with high PPT scores tended to display reduced CKC compared to what is expected from their BBT score (below regression line). The same conclusions can be drawn when standardized BBT scores are subtracted from standardized CKC, and this contrast is plotted against PPT scores (Fig. 5C). Indeed, when PPT scores are low, the contrast was high, meaning that a participant had a CKC value that was higher than expected from their BBT score, and vice versa, as attested by a significant negative correlation (*r* = −0.35, *p* = 0.006).

## Discussion

In this study, we examined the relationship between CKC, a marker of cortical proprioceptive processing, and motor performance, with the aim of establishing the behavioral relevance of CKC. We identified a significant relationship between CKC at F1, but not F0, and motor performance of a gross motor control task. Conversely, the motor performance of a fine motor control task was informative about CKC, but only when considered in synergy with gross motor ability. Finally, motor coordination did not contribute to motor performance, but showed a trend of association with CKC at F0, but not F1.

### Behavioral relevance of CKC

In our study, higher BBT scores, indicating more favorable gross motor function, corresponded to stronger CKC. This finding suggests that there is a behavioral advantage to stronger CKC on this motor task. In the following paragraphs, we present several non-exclusive explanations for this behavioral advantage.

Previous analyses have shown that CKC likely reflects the processing of afferent somatosensory, mostly proprioceptive, information (Piitulainen et al., 2013a; Bourguignon et al., 2015). Therefore, increased CKC could indicate an increased quantity of cortical resources allocated for sensory processing. Following the views whereby inefficient neural coding could be compensated by recruiting additional neural populations (Nurmi et al., 2021), stronger CKC may represent an increase in the utilization of neural substrate. This idea has been brought forward in another study in which stronger CKC was associated with worse postural stability (Piitulainen et al., 2018b). But unlike during postural balance, where the consistent need for more neural resources may be computationally unfavorable, the ability to allocate additional neural substrates for a novel task, like the BBT in our study, could foster better integration of afferent signals. Accordingly, interpretation of CKC magnitude appears to be task–dependent.

The relationship between CKC and task performance might also be explained by a more faithful cortical representation of proprioceptive afferences in the best performers. This is well in line with the behavioral relevance of neural entrainment to external stimuli seen across a range of modalities, best explored in language literature where entrainment to speech benefits comprehension (Ahissar et al., 2001; Peelle et al., 2013; Vanthornhout et al., 2018; Bertels et al., 2023), as well as in the context of sign language observation (Malaia et al., 2021). Alternatively, higher BBT scores are likely to be achieved with faster movements. Since muscle spindle activity has been shown to increase with faster movement (Sasaki et al., 2018), it should result in stronger afferent volleys to the SM1, and hence increased CKC. Attention and motivation may also vary across participants and impact both performance and CKC through the neuronal resources allocated to the task. Thus, changes in CKC, similar to corticomuscular coherence, could be related to attention-based parameters (Kristeva-Feige et al., 2002; Johnson et al., 2011).

Beyond the behavioral relevance of CKC for gross motor function, our PID analysis revealed that fine motricity, in the form of PPT scores, brought more intricate synergistic information about CKC when observed together with BBT scores. We show that individuals with enhanced fine motor skills (high PPT) tend to possess lower levels of CKC than expected, based on their gross motor skills (BBT), with the converse also holding true. This suggests that CKC is higher in relation to features present in the BBT, but not in PPT. While both tasks rely on gross control of the arm and forearm, performance of the BBT likely depends on accurate proprioceptive encoding of upper limb joint positions and movements, while the PPT may rely more on visuomotor integration as required by the precise placement of the peg (Hinkle and Pontone, 2021), giving less weight to proprioception. These views lay strong support to the notion that CKC is mainly driven by proprioceptive afferences (Piitulainen et al., 2013a; Bourguignon et al., 2015), while adding the important notion of behavioral relevance.

In assessing the relationship between CKC and BBT values, we identified the need to correct CKC values for EEG SNR, leading to a slight decrease in correlation strength from 0.41 without correction to 0.29 with correction. However, this drop does not necessarily indicate that the initial association between non-corrected CKC values and BBT scores was inflated. Indeed, correcting CKC values for EEG SNR was meant to control for a bias, but also removed part of the effect we meant to identify. Regarding the bias, it could arise because the EEG noise level typically decreases with the frequency (Demanuele et al., 2007), leading to higher CKC values at higher frequencies if the amplitude of the genuine cortical signal is constant. However, a previous study, though conducted on a smaller sample size than the present one, reported that CKC was not affected by movement frequency within the range 1–3 Hz (Marty et al., 2015b). Regarding the second point, our hypothesis was that of an association between CKC and motor performance, which in the case of BBT is proportional to movement frequency. Yet, the coherence is well known to increase with genuine signal amplitude, along with better phase locking. Hence, under our hypothesis, it was expected that EEG SNR would increase with motor performance. Therefore, the correlation of 0.29 obtained with corrected CKC values should be taken as a lower bound for the strength of the association between CKC and motor performance.

While we cannot argue that motor performance depends on genetic factors giving rise to increased or decreased sensory processing ability, or occurs as a result of a lifetime of motor practice, there is evidence for both to be true. Recently, the *Piezo2* gene, expressed primarily in peripheral sensory neurons, but also found in the brain, has been linked to interindividual variability in the functionality of mechanosensation (Nagel and Chesler, 2022), suggesting the efficacy of proprioceptive processing could be dependent on genetics. Alternatively, motor skill practice and motor learning are two extremely well described behaviors known to modify brain activity (Kantak and Winstein, 2012).

### Coordination

In using EMG recordings during the BBT, we assessed muscle coordination and identified how it relates to motor performance and CKC. Here, coordination refers to the similarity in muscle activation patterns across movement cycles.

We did not identify a significant relationship between muscle coordination and BBT scores. Although we expected that increased coordination values would coincide with higher BBT scores, our result is not entirely surprising. In the BBT, blocks sit randomly within one partition. Participants must adjust each block transfer depending on their selection, resulting in variable joint trajectories (Hebert et al., 2014) and correspondingly variable muscle activation patterns. Therefore, a flexible motor control strategy in which muscle activation patterns differ depending on block selection may point towards efficient motor control, while coordination, as defined in this study, is less relevant.

Correlations for CKC at F0 and F1 and coordination revealed no significant relationships, except for a weak one when CKC at F0 was corrected for EEG SNR. These findings support our initial hypothesis that CKC and coordination hinge on different mechanisms.

### Limitations and perspectives

Owing to the dynamic nature of the task, our EEG recordings were contaminated to some degree with movement artifacts. During the BBT, participants were instructed to keep their heads still and facial muscles relaxed, but in practice, this was challenging. This likely reduced the precision of CKC estimation, and could explain the lack of relationship identified between motor performance and CKC at F0. Artifacts were noticeable as increased coherence at electrodes on the edge of topographic plots, which is why these were excluded in the analyses.

Another reason why CKC at F0 was not found to be behaviorally relevant could rest in its susceptibility to heartbeats and resultant artifacts, which tend to occur around the same frequency. However, heartbeat artifacts were sought and suppressed with ICA. Alternatively, previous studies have also suggested that CKC at F1 may not merely represent the harmonic of CKC at F0 (Pollok et al., 2004, 2005; Bourguignon et al., 2012; Piitulainen et al., 2013a). For simple flexion/extension movements, it could arise due to the integration of the two volleys of sensory information caused by flexion and the extension within a movement cycle (Pollok et al., 2004, 2005; Bourguignon et al., 2012; Piitulainen et al., 2013a). Likewise, during the BBT, participants performed back and forth movements during one cycle.

The present study opens up important perspectives for future clinical studies and practice. That is, CKC and coordination represent informative measures that could be gathered in the framework of the BBT, providing a more global view of a patient’s motor control, and hopefully improve clinical decision making.

### Conclusion

Our study confirms the ability to evaluate CKC during a dynamic and variable task such as the BBT and establishes CKC as a behaviorally relevant marker of effective cortical proprioceptive processing, where high values of CKC are predictive of good gross motor function. CKC measurement may favorably complement classical behavioral assessments of proprioception in contexts where proprioceptive acuity is in question, help discern if the brain is involved in proprioceptive deficits, or assess plasticity related to improving cortical processing.

## Acknowledgments

Scott Mongold was supported by an Aspirant research fellowship awarded by the F.R.S.-FNRS (F.R.S.-FNRS, Brussels, Belgium; grant FC 46249). Christian Georgiev was supported by an Aspirant research fellowship awarded by the F.R.S.-FNRS (F.R.S.-FNRS, Brussels, Belgium; grant 1.A.211.24F). Scott Mongold, Thomas Legrand, and Mathieu Bourguignon were supported by the Fonds de la Recherche Scientifique (F.R.S.-FNRS, Brussels, Belgium; grant MIS F.4504.21).

